# Tree bark as a substrate for mycelium-bound composites with two *Ganoderma* species

**DOI:** 10.1101/2025.06.16.659851

**Authors:** Tim K. Felle, Jannis Estenfelder, Charlett Wenig, Ferréol Berendt, Michaela Eder, Tian Cheng, J. Philipp Benz

## Abstract

Bark is currently considered as a by-product of the wood industry and is mostly incinerated for energy, left in forests, or used as a mulch layer in gardens, parks, and forests to help prevent soil from drying out. However, considering that bark makes up about 10-20% of the tree volume, there is a considerable amount of material that can be exploited and should be investigated in terms of a resource-efficient bio-economy. One way to use bark as a raw material for innovative products could be as a substrate within mycelium-bound composite materials. However, since one of the natural functions of bark is to inhibit microbial infestation of the trees, it was unclear whether bark could be utilized in this manner. Therefore, we investigate in this study the possibilities of producing such composites by evaluating the performance of several bark-fungus combinations. Three different barks (from Douglas fir, Scots pine and European birch) and two species of fungi (*Ganoderma sessile* and *Ganoderma adspersum*) were selected for the experiments. Mycelium growth rates were evaluated with a newly developed method using fungal “growth tubes”. In addition, composites were prepared for performance tests from pure bark and 1:1 mixtures of bark and beech wood sawdust. Composites made of mixed bark and beech wood were mostly well overgrown with a thick layer of mycelium on the surface, supporting higher compression strengths. The mycelium layer on the composites made with tree bark only was considerably thinner, resulting in lower compression strength. Water absorption potential was found to be highly dependent on the mycelium layer on the composite surfaces, which has substantial hydrophobic properties. Overall, although the required incubation times tend to be longer than for other commonly used substrates, our experiments demonstrate that bark clearly represents a potential co-substrate for the production of mycelium-bound composites. Nevertheless, further studies with different fungi and barks are required to identify mixtures for specific applications that require shorter incubation times.

## Introduction

With growing awareness of the finite availability of fossil-based resources, their environmental impact, and the urgent need for more sustainable production and consumption models, the bio-based circular economy is expected to expand in the coming years [1]. This transition necessitates the efficient utilization of industrial side streams and waste materials, converting them into biodegradable products to support sustainable resource management and waste reduction [2]. As a renewable and sustainable resource, wood plays a vital role in the circular economy, offering an environmentally friendly alternative to conventional materials, particularly in the construction and housing industries [3]. The European construction sector, for example, is a major consumer of resources, accounting for approximately 50% of extracted materials and energy use, as well as one-third of water consumption and waste generation [4]. However, woodworking processes inherently generate substantial by-products and residual materials, which contribute significantly to industrial wood waste [5]. During primary wood processing, logs yield various by-products, including sawdust, wood chips, wood particles, and bark [6], presenting both a challenge and an opportunity for sustainable material utilization. While bark has economic and functional potential, including applications in agriculture, medicine, and panel manufacturing [7–9], most of it is either disposed of in landfills or incinerated for energy production [10]. Bark accounts for approximately 10-20% of a tree’s total volume, generating over 20 million tons annually in the United States and between 6-8 million tons in Europe [11, 12]. Despite this huge mass, bark remains an underutilized resource - primarily due to its structural complexity and variability within and between tree species. Developing innovative cascading utilization strategies for bark could contribute to more sustainable and value-added applications.

Bark is compositionally distinct from wood, consisting of lignin (38-55%), polysaccharides (30-65%), and a significant fraction of extractives, which range from 2-35% in coniferous trees and 5-15% in deciduous species [13, 14]. Among these extractives, polyphenols, including tannins, play a critical role in the tree’s natural protective mechanisms due to their antifungal, antimicrobial, and antioxidant properties [15–18]. Additionally, bark contains a periderm layer with cork, which consists of lignin, polysaccharides, and substantial amounts of suberin, a hydrophobic biopolymer known for its antimicrobial properties [19]. The suberin content in bark varies significantly across species, with European birch and Douglas fir, for example, containing over 30 wt.% [20], while Scots pine bark contains only about up to 2% [21]. In addition, birch bark also contains betulin, a pentacyclic triterpene alcohol with strong antimicrobial properties [22]. Given these chemical protective mechanisms, it remained unclear whether fungi can effectively colonize bark to form mycelium-bound composites. Fungi have the ability to convert organic materials into a diverse range of valuable products [23], making them an essential tool in sustainable biotechnology. Mycelium-bound composites are composed of renewable organic substrates bonded by fungal mycelium, providing complete biodegradability at the end of their lifecycle [24]. These materials are extensively studied as eco-friendly alternatives for applications in packaging, insulation, and lightweight construction [25, 26]. Identifying suitable wood-inhabiting fungi from the phylum Basidiomycota [27] is key to optimizing substrate compatibility and composite performance.

Given the abundant availability of bark and its potential as a material source for the bioeconomy, this study aimed to assess its feasibility as a substrate for mycelium-bound composites by evaluating fungal colonization, composite properties, and material performance. The bark of three tree species (*Pseudotsuga menziesii*, *Pinus sylvestris*, and *Betula pendula*) was tested with two wood-inhabiting fungal species (*Ganoderma sessile* and *Ganoderma adspersum*). Growth experiments, material performance tests, and morphological analyses were conducted to examine mycelium colonization, density, mechanical strength, and water absorption properties, providing insights into the suitability of bark as a substrate for mycelium-bound composite materials.

## Material and methods

### Fungal species

In this study, two *Ganoderma* species, *Ganoderma adspersum* and *Ganoderma sessile*, were used. The wild-type strain of *G. sessile* was a generous gift from Harald Kellner (Technical University of Dresden – IHI Zittau) and was originally isolated from a fruiting body collected in Kentucky (USA). A fruiting body of *G. adspersum* was collected from a living maple tree in the city park of Freising (Germany). Small tissue samples (∼1.5 mm³ each) were excised and immersed in a 10% H_₂_O_₂_ solution for 3 minutes to minimize bacterial and fungal contamination. The disinfected samples were transferred onto sterile potato dextrose agar (PDA) plates (39 g PDA per 1 L ddH_₂_O; Sigma-Aldrich, St. Louis, USA) under sterile conditions. The plates were incubated at 26 °C and 90% relative humidity (RH) under reduced light for 48 hours. The resulting vegetative mycelium was subcultured onto fresh PDA plates and incubated for an additional 7 days. Strain identity was confirmed by Sanger sequencing of the internal transcribed spacer (ITS) region using primers ITS1 (TCCGTAGGTGAACCTGCGG) and ITS4 (TCCTCCGCTTATTGATATGC) [28]; of the translation elongation factor 1-alpha (TEF1) gene using primers TEF1-983F (GCYCCYGGHCAYCGTGAYTTYA) and TEF1-1567R (ACHGTRCCRATACCACCSATCT) [29]; and of the RNA polymerase II subunit 2 gene (rpb2) using primers bRPB2-6F (TGGGGYATGGTNTGYCCYGC) and bRPB2-7R (GAYTGRTTRTGRTCRGGGAAVGG) [30].

### Substrate preparation and characterization

Three different types of tree bark were used for the experiments in this study, i.e. bark from Scots pine (*Pinus sylvestris*), European birch (*Betula pendula*), and bark from Douglas fir (*Pseudotsuga menziesii*). Bark was ground to a particle size of 0,5 - 7.0 mm using a cutting mill (Retsch, Haan, Germany). Beech (*Fagus sylvatica*) wood sawdust (particle size 0.5 – 1.0 mm) (J. Rettenmaier & Söhne GmbH + Co KG, Rosenberg, Germany) was used as a reference substrate. Prior to the experiments, all substrates were dried in an oven at 103°C for 24 hours to accurately determine the dry weight for further experiments. In addition to the preparation of pure bark, 50:50 bark-to-beech wood mixtures (by weight) were also prepared (Fig. 1; Tab. 1). The pH value of all substrates was measured by adding 4.0 g of each substrate to 50 mL of ddH_2_O and continuously shaking the mixture at 170 rpm for 12 hours. The pH value was then measured with a pH meter (SevenEasy^TM^, Mettler Toledo International Inc., Columbus, USA).

**Fig. 1:**
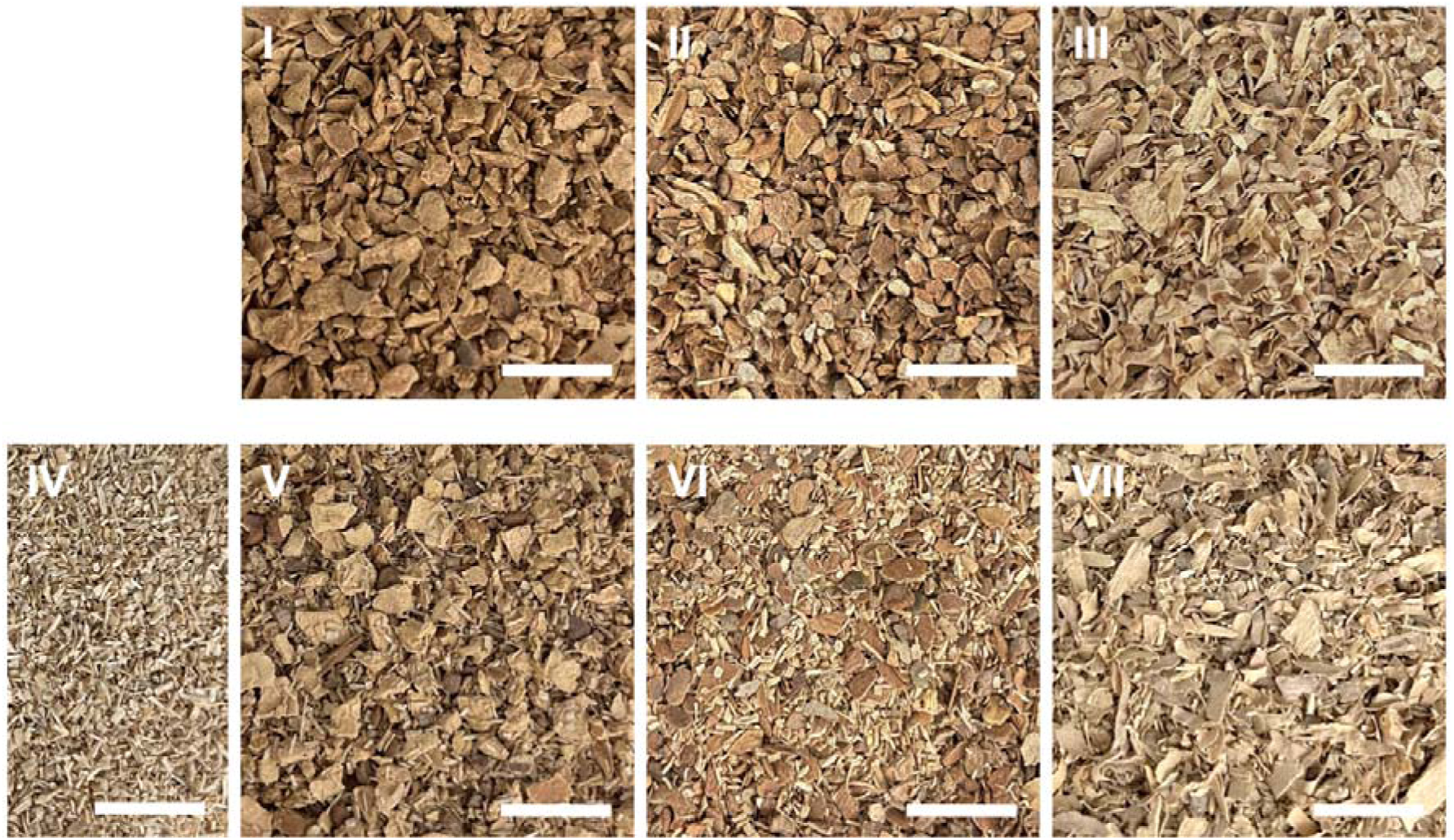
Overview of the substrates used in this study. The study included three types of pure bark: Douglas fir (I), Scots pine (II), and European birch (III), along with European beech wood sawdust (IV) as a reference material. In addition, three mixed substrates were prepared, each consisting of a 50:50 ratio of bark to beech wood sawdust: Douglas fir bark with beech wood (V), Scots pine bark with beech wood (VI), and European birch bark with beech wood (VII). Scale bars represent 10 mm.

**Tab. 1:**
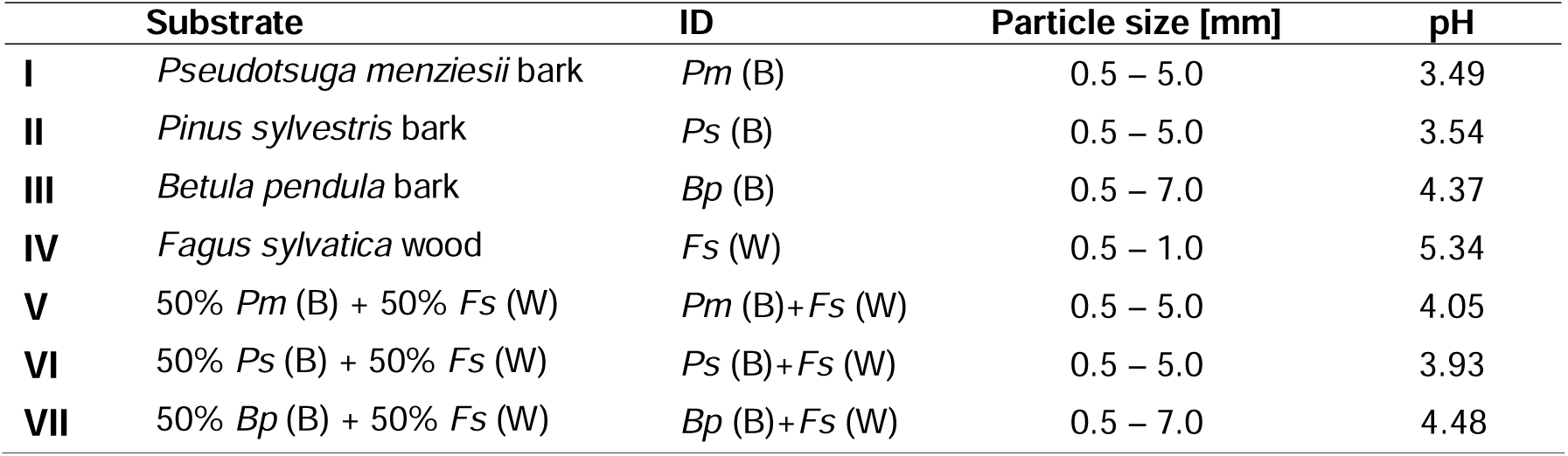
Substrates used in this study with abbreviations (ID), particle sizes, and pH values.

### Media and pre-cultures preparation

All media were sterilized by autoclaving at 121°C for 20 minutes prior to utilization. For the preparation of liquid cultures, 100-mL flasks were filled with 50 mL of potato dextrose yeast (PDY), consisting of 1 L of ddH_2_O, 24 g of potato extract glucose broth (Carl Roth GmbH + Co. KG, Karlsruhe, Germany), and 5 g of yeast extract (Sigma-Aldrich, St. Louis, USA). Following sterilization in an autoclave (Systec VX-150, Systec GmbH & Co. KG, Linden, Germany), the flasks were cooled to room temperature and subsequently filled with five agar plugs (Ø 8 mm) containing fungal mycelium obtained from the periphery of Petri dishes filled with potato dextrose agar (PDA). These Petri dishes were stored in a cooling room at 4°C. The flasks were then covered with aluminum lids and placed into an incubator shaker (New Brunswick Innova® 42, Eppendorf SE, Hamburg, Germany) set to 26°C and a shaking speed of 100 rpm. The samples were left to incubate in this environment for a period of seven days.

After 5 days of incubation, fungal biomass had spread throughout the PDY in the form of mycelium balls. The residual media was then decanted, and the fungal biomass was homogenized for approximately 10 seconds using an Ultra-Turrax® dispersing instrument (TP 18/10, IKA®-Werke GmbH & Co. KG, Staufen, Germany). To prepare the grain spawn, the mycelium suspension from a single flask was transferred into a jar containing sterilized rye grains. The rye grains were prepared by placing 100 g of grains with 150 mL of ddH_₂_O in a 580-mL Weck glass jar (Weck glass and packaging GmbH, Bonn, Germany) and soaking them for 1 hour at 60°C in an oven. After soaking, the excess water was decanted, 3 g of PDB was added, and the mixture was autoclaved. Once the mycelium suspension was incorporated, the grain spawn jars were sealed with medical tape and placed in an incubator under controlled conditions (26°C, 90% RH) with reduced light for 5–7 days. After incubation, the grains were sufficiently colonized with mycelium, making the grain spawn ready for further applications.

### Growth rate assessment

The fungal growth rate on different substrates was assessed using a newly developed fungal “growth tubes” method (Fig. 2). The “growth tubes” were prepared as follows: 15-mL laboratory falcons (Sarstedt AG & Co. KG, Nümbrecht, Germany) were cut at the top and sealed with medical tape to ensure airflow. Subsequently, the growth tubes were filled with 4 g of each substrate and 6 mL of ddH_2_O, and then autoclaved. Following the autoclaving process, the growth tubes were cooled, opened in the screw area, and inoculated with 10 pieces of rye grains from the grain spawn. This inoculation was performed under sterile conditions using laboratory tweezers. The tubes were then sealed and placed into the incubator, which was set to a reduced light and controlled climate (26°C and 90% RH). For each substrate-fungus combination eight biological replicates were prepared. The length of the mycelium was measured at four points on the growth tube section at 24-hour intervals. The mean daily value of these four measurements was expressed as the growth rate in millimeters per day. Upon reaching the maximum line of the growth tube, the tubes were cut, and the overgrown substrate was removed. The mycelium growth was then also visually assessed.

**Fig. 2:**
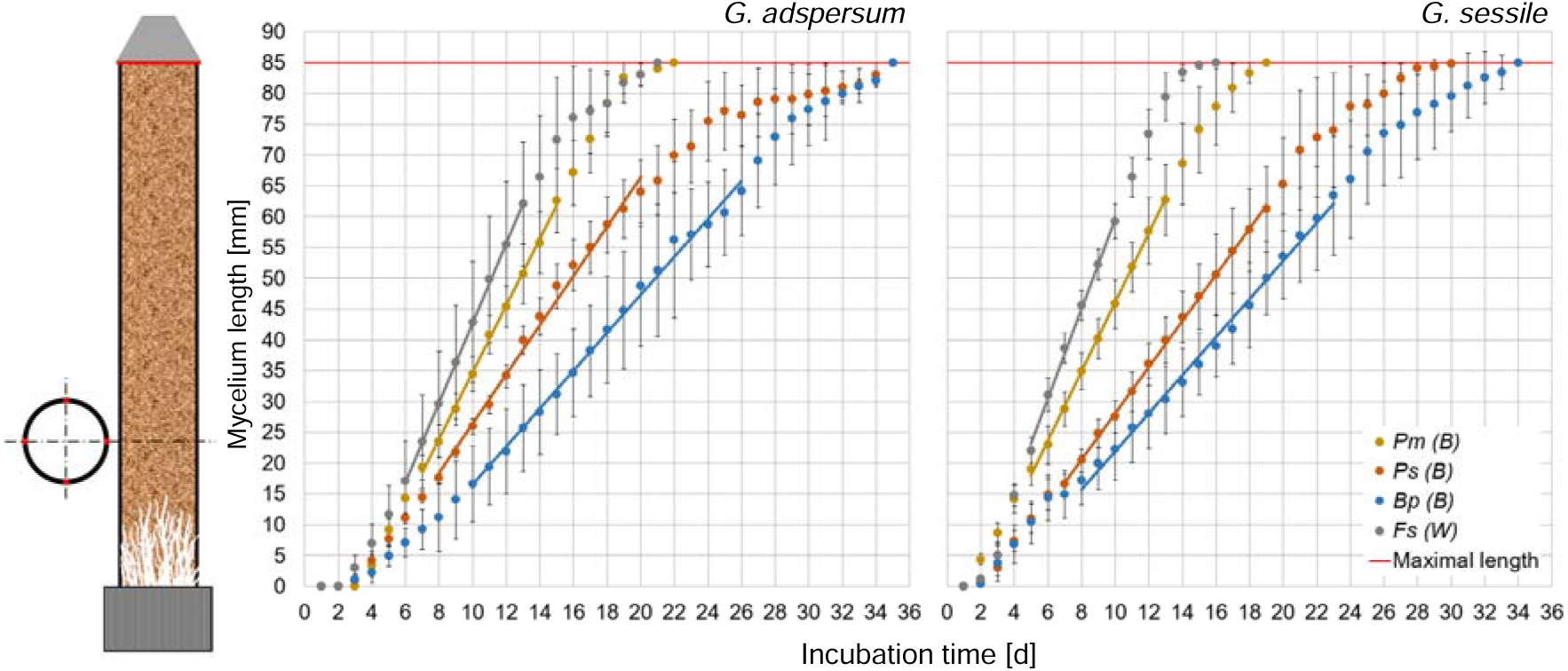
Growth tube setup and mycelium growth [mm] over time [d] for *G. adspersum* (left) and *G. sessile* (right) on different substrates: *Pm* (B) – Douglas fir bark; *Ps* (B) – Scots pine bark; *Bp* (B) – European birch bark; *Fs* (W) – European beech wood. The red line indicates the maximum possible growth length. Error bars represent standard deviations (n = 8).

### Fabrication of composites

Plastic bags with a grid for air exchange (Prolinx, Düsseldorf, Germany) were filled with 100 g substrate, 125 mL ddH_2_O, and 3 g potato extract glucose broth, and then autoclaved. All the substrates were inoculated with three different amounts of grain spawn (10, 20, and 30%). Prior to inoculation, the grain spawn was milled in a blender (Bosch CNSM20, BSH Hausgeräte GmbH, München, Germany) for approximately 20 seconds and then weighed using a precision balance (PLJ 600-3CM, KERN & SOHN GmbH, Balingen-Frommern, Germany) under sterile conditions. Subsequent to cooling, the predetermined quantity of grain spawn was added to the plastic bag, thoroughly mixed with the substrate, and sealed with tape. Subsequently, the substrates were placed into the incubator, which was set to a reduced light and controlled climate (26°C and 90% RH) for approximately 8-12 days. The fabrication of composites was only possible when the substrates, which were enclosed in plastic bags, had undergone sufficient overgrowth, as indicated by the presence of a white surface on the substrate. The overgrown substrate was then loosened by hand under sterile conditions and placed into two different cylindrical molds, depending on the subsequent test. The filled molds were subsequently transferred into a storage box (SAMLA, Inter IKEA Systems B.V., Delft, Netherlands) and incubated at 26°C and 90% RH for 3-4 days. Following this, the molds were removed under sterile conditions, and the samples were then again subjected to a 2-3 days incubation period. Afterwards, the composites were retrieved from the incubator, measured with a caliper, and dried at 70°C for 24 hours. Following this, the composites were measured once more to determine their final dimensions.

For the compression tests, triplicate specimens measuring 27 mm × 70 mm (Ø × height) were prepared for all substrate combinations. For the water absorption potential test, specimens measuring 86 mm × 13 mm (Ø × height) were prepared (n = 4), but only using substrate combinations containing 10% grain spawn. Additionally, some specimens with the same dimensions as those used in the water absorption test were prepared for morphological analysis.

### Density measurements

Density was determined only for the specimens used in the compression tests. All specimens were cut to a uniform length of 60 mm using a circular saw. Following this process, the composites were conditioned under controlled laboratory conditions (21°C and 60 % RH), then measured and weighed to calculate their density using the formula ρ = m/V [kg/m³], where m represents the specimen’s weight [kg] and V denotes its volume [m³].

### Compression test

The compression test was conducted using a universal testing machine (1455 Retro Line, ZwickRoell GmbH & Co. KG, Ulm, Germany) with a crosshead speed of 10 mm/min and a preload of 1 N. The test was automatically terminated when the force dropped by more than 5% within 0.1 seconds. Force and displacement data were recorded every 0.1 seconds using TestXpert III software (ZwickRoell GmbH & Co. KG, Ulm, Germany). Engineering stress (σ) was calculated as the applied force (F) [N] divided by the initial cross-sectional area (A) [mm²] of the specimens (σ = F/A [MPa]). Strain (ε) was determined using the formula ε = ΔL/L_₀_, where L_₀_ [mm] is the original height of the specimen. Based on the resulting stress–strain diagrams, the modulus of elasticity (E) was calculated in the linear elastic region using the relationship E = σ/ε [MPa]. The ultimate compressive strength was defined as σ_max_ = F_max_/A [MPa], where F_max_ is the maximum force recorded during the compression test.

### Water absorption test

This test was conducted in accordance to the respective standard [31], describing the determination of water absorption coefficient by partial immersion. Prior to testing, the prepared composites were dried at 60°C for 12 hours and sealed on the sides with waterproof Parafilm® (M Laboratory Film, Bemis Inc., Neenah, USA) to ensure that only the bottom surface was in contact with water. Each specimen was weighed to the nearest 0.01 g using a precision balance before immersion. For the water container, a storage box (39 × 28 × 14 cm³) was used. Specimens were placed on plastic rings (4 mm in height, 37 mm in diameter) to keep them elevated above the container’s base. To maintain stability during immersion, each specimen was weighted with 50-mL laboratory conical tubes filled with sand, with an average mass of 63 g. The container was then carefully filled with 600 mL of demineralized water, ensuring that the specimens were half-immersed, and the timing of the experiment commenced. At predefined intervals, the specimens were removed, briefly placed on a paper towel to eliminate excess surface water, weighed to the nearest 0.01 g, and then immediately re-immersed. To maintain consistent test conditions, the water level in the container was regularly replenished. The experiment was conducted for 72 hours, with more frequent measurements taken within the first 12 hours.

### Morphological analysis

The morphology of fungal growth on all substrate combinations was initially analyzed visually using a binocular microscope (S8 APO, Leica Microsystems GmbH, Wetzlar, Germany). To investigate hyphal growth at high resolution, samples with a size of 1.3 × 2 × 2 cm³ were cut from the central region of the specimens and sputter-coated with a 10 nm platinum layer at 30 mA using a CCU-010 coating unit (Safematic GmbH, Zizers, Switzerland). Afterwards the samples were transferred to an Environmental Scanning Electron Microscope (ESEM Quattro S, Thermo Fisher Scientific, Brno, Czech Republic) operated in the high vacuum mode with an acceleration voltage of 5 keV.

### Data processing and statistical analysis

Growth kinetics were evaluated by calculating mean values and standard deviations to assess variability. Growth rates were analyzed using trendlines and the corresponding coefficients of determination (R²), calculated in Microsoft Excel (Microsoft Corporation, USA). Stress–strain curves were analyzed using OriginPro (OriginLab Corporation, USA), with the slopes determined by linear fits. Due to the time-consuming sample preparation, the number of replicates per sample group was limited to three; hence, no averages or standard deviations were calculated for these data.

### Results and Discussion Growth of mycelium on bark

Bark exhibits a complex anatomical structure that reflects its diverse functions in plants and the secondary growth of trees. It consists of inner bark (secondary phloem, cortex, phelloderm) and outer bark (phellem or rhytidome) [32]. This structural complexity and the significant chemical variation between layers might affect fungal colonization. Therefore, the “growth tube” method was developed to partially homogenize the substrate by milling it into smaller, mixed particles, allowing for a more quantitative assessment of fungal growth. Distinct mycelium growth patterns were observed, with both fungal species exhibiting similar trends (Fig. 2). Both *Ganoderma sessile* and *Ganoderma adspersum* exhibited the fastest growth on the reference substrate, beech wood. *G. sessile* reached the maximum tube length of 85 mm in 16 days, while *G. adspersum* took 21 days. On the other hand, both fungi showed slightly slower growth on Douglas fir bark, with *G. sessile* completing its growth in 19 days and *G. adspersum* in 22 days. Even slower growth rates were observed on Scots pine and European birch bark. *G. sessile* reached the maximum on pine bark in 30 days and *G. adspersum* in 35 days. For birch bark, *G. sessile* reached maximum growth in 34 days, while *G. adspersum* also required 35 days. In contrast to wood, which is primarily composed of polysaccharides and lignin, bark contains a significantly higher percentage of polyphenols (extractives) and suberin [14]. These compounds are highly resistant to enzymatic degradation due to their hydrophobic properties and complex structures, which might contribute to the slower growth of both fungal species on bark substrates. Additionally, birch bark is particularly rich in betulin, a natural triterpenoid known for its strong antibacterial and antifungal properties [33].

The lag phase duration varied between fungal species and substrate types. The longest lag phase was observed for *G. adspersum* on Doulgas fir bark (3 days), whereas *G. sessile* showed initial growth by day 2 on all substrates. Following the lag phase, all fungi entered an accelerated growth phase, where the mycelium – now in direct contact with the substrate - likely began utilizing its nutrients for sustained growth [34, 35]. Subsequently, this phase was marked by an even spread of mycelium across the tube. The linear growth region, spanning from 15 mm to 65 mm, was selected for calculating growth rate (mm d^-1^) and coefficient of determination (R²; Tab. 2). Growth within this range was highly correlated for all combinations, indicating that the fungi exhibited a consistent growth pattern once the initial lag phase was overcome. This also supports the effectiveness of the growth tube method in partially homogenizing the substrate through milling, thereby reducing the impact of structural and chemical variations on fungal colonization. Interestingly, no significant differences in growth rates were observed between the two fungal species in the linear growth region, likely due to their shared genus. However, for both species— more pronounced in *G. adspersum*—the growth rate declined after reaching approximately 75 mm in the growth tube. This decline may be attributed to the increased oxygen availability near the open end of the tube, leading to denser mycelial growth within the substrate, which is not visible on the tube’s surface.

**Tab. 2:**
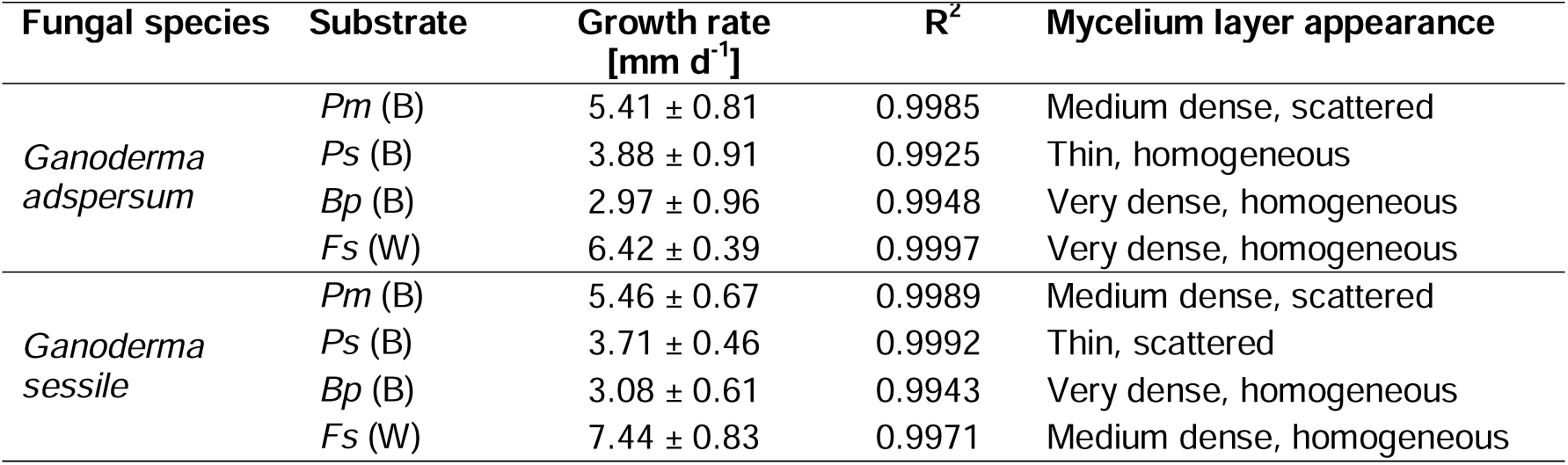
The mycelium growth rate (mm d^-1^) on bark substrates, including the coefficient of determination (R²) to assess growth consistency, is analyzed within the linear growth region ranging from 15 mm to 65 mm. Additionally, macroscopic traits of the mycelium layer are examined for each substrate.

| Fungal species | Substrate | Growth rate<br>[$\text{mm d}^{-1}$] | $R^2$ | Mycelium layer appearance |
| --- | --- | --- | --- | --- |
| *Ganoderma adspersum* | *Pm* (B) | $5.41 \pm 0.81$ | 0.9985 | Medium dense, scattered |
| | *Ps* (B) | $3.88 \pm 0.91$ | 0.9925 | Thin, homogeneous |
| | *Bp* (B) | $2.97 \pm 0.96$ | 0.9948 | Very dense, homogeneous |
| | *Fs* (W) | $6.42 \pm 0.39$ | 0.9997 | Very dense, homogeneous |
| *Ganoderma sessile* | *Pm* (B) | $5.46 \pm 0.67$ | 0.9989 | Medium dense, scattered |
| | *Ps* (B) | $3.71 \pm 0.46$ | 0.9992 | Thin, scattered |
| | *Bp* (B) | $3.08 \pm 0.61$ | 0.9943 | Very dense, homogeneous |
| | *Fs* (W) | $7.44 \pm 0.83$ | 0.9971 | Medium dense, homogeneous |

Conversely, prolonged exposure to air at the open end may have caused localized drying of the substrate, thereby limiting water availability and reducing fungal growth kinetics. Further experiments using growth tubes of varying lengths and different opening designs to control air circulation are needed to clarify the factors contributing to this decline in fungal growth rate. Once the mycelium reached its maximum growth in the tubes, the overgrown substrate was removed, and the density and growth characteristics of the mycelium were visually assessed (Tab. 2). Interestingly, despite the differences in growth rates, the mycelium on birch bark (which showed the slowest growth) and beech wood (which showed the fastest growth) was remarkably similar. This suggests the presence of potential metabolic or morphological adaptations that may allow the fungi to optimize nutrient absorption, even under less favorable conditions. This could also be related to the pH values of the substrates (Tab. 1). Notably, Douglas fir bark, with the lowest pH (3.49), supported the fastest mycelial growth but exhibited a moderately dense and scattered mycelium layer. In contrast, birch bark, with the highest pH (4.37) among the bark substrates, was associated with the slowest growth, though the mycelium of both fungi grew very densely and completely. Previous studies [35, 36] have shown that *Ganoderma* spp. can grow within a broad pH range of 5 to 9. The faster growth observed at lower pH values in the present study may be attributed to a mechanism described here as ‘escape growth’, where the mycelium accelerates through unfavorable conditions in search of a more suitable environment. This phenomenon may also be linked to the macroscopic characteristics of the mycelium, as substrates with lower pH exhibited a thinner and more scattered mycelium layer.

### Density

The density of the cylindrical mycelium composites used in the compression tests was calculated to assess its variation in relation to the three key variables examined in this study: substrate composition, fungal species, and the amount of grain spawn (inoculum). The densities ranged from 0.26 g/cm³ for beech wood to 0.57 g/cm³ for birch bark in composites made with *G. adspersum*. Composites prepared with *G. sessile* generally exhibited higher densities, ranging from 0.31 g/cm³ for beech wood to 0.59 g/cm³ for birch bark. In most cases, composites derived from mixed substrates (50% bark and 50% wood) exhibited lower densities than those composed solely of bark. This reduction in density is likely attributable to the lower density of wood particles as well as to a thicker mycelium layer on the surface. Composites with mixed substrates inoculated with *G. sessile* exhibited densities ranging from 0.41 g/cm³ (Douglas fir bark with beech wood) to 0.51 g/cm³ (birch bark with beech wood). In contrast, mixed-substrate composites prepared with *G. adspersum* showed a broader density range, with Douglas fir bark and beech wood yielding 0.28 g/cm³ and birch bark with beech wood yielding 0.47 g/cm³. Overall, higher grain spawn concentrations led to increased composite densities, as rye grains exhibit a dry density of approximately 0.70 g/cm³. However, no relationship was observed between the quantity of grain spawn and the thickness of the mycelium layer.

### Mechanical properties

Three biological replicates were prepared and tested in compression for each combination of substrate, fungal species, and grain spawn percentage (126 samples in total). Stress–strain curves were plotted to investigate the compression properties. Two distinct types of stress–strain curves were observed (Fig. 3a). The majority of the curves exhibited a steeper initial slope, while some samples showed a lower initial slope that increased gradually during the test. Due to this gradual increase, issues with sample or machine misalignment are highly unlikely. The two types of stress–strain curves were analyzed as follows: (A) slope of the initial linear region; (B) slope of the second linear region; and (C) ultimate compressive strength.

**Fig. 3:**
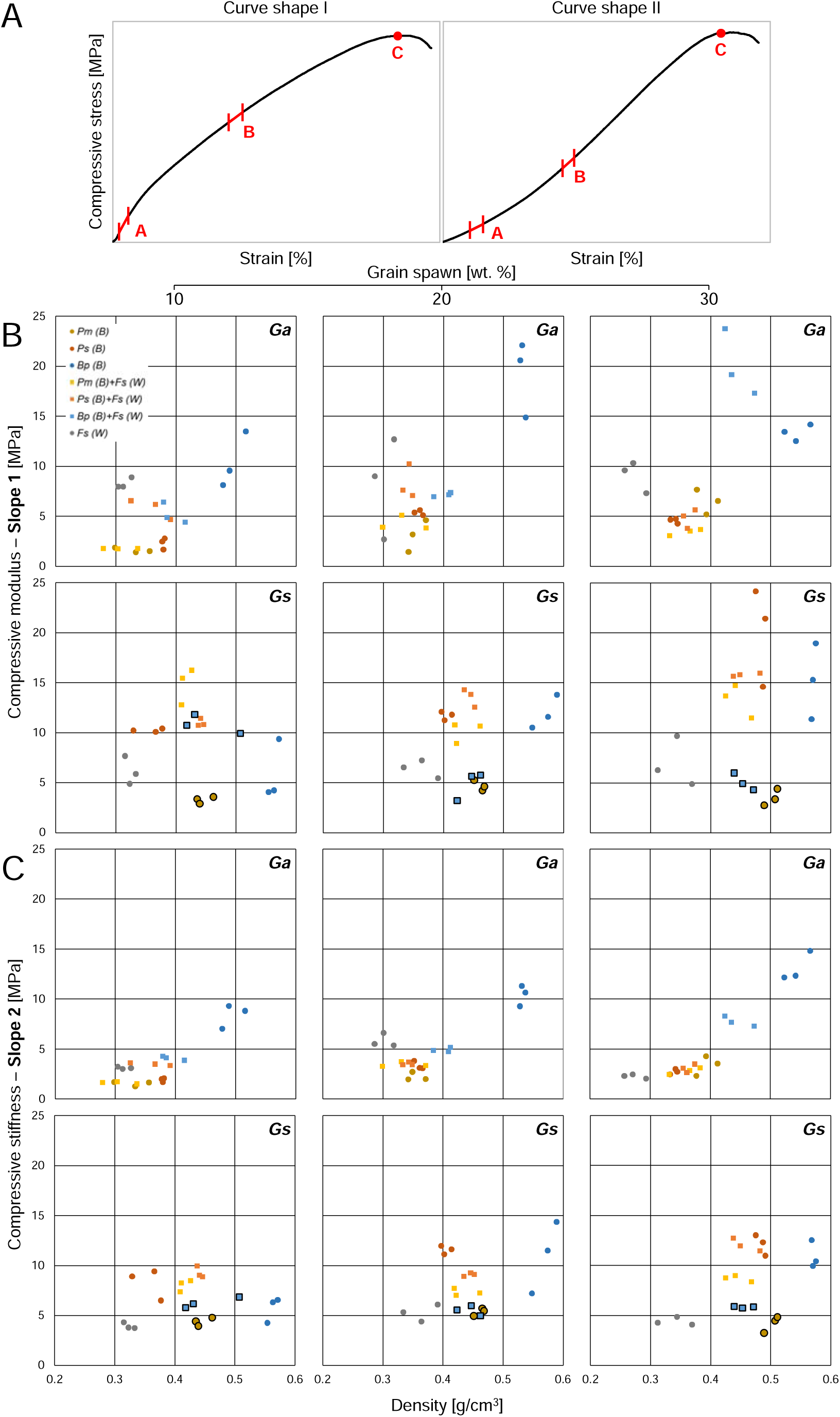
**A** Representative compressive stress–strain curves illustrating: (A) the initial linear region (Slope 1, elastic response), (B) the second linear region (Slope 2, associated with densification or stiffening), and (C) the ultimate compressive strength (maximum stress before failure). Curve shape II was observed in composites made from Douglas fir bark (*Pm (B)*) and from a mixture of birch bark and beech wood (*Bp (B)* + *Fs (W)*) inoculated with *G. sessile*. **B** Compressive modulus values derived from the initial linear region A (Slope 1) plotted against density. **C** Compressive stiffness values derived from the second linear region B (Slope 2) plotted against density. Data are shown separately for each grain spawn level (10, 20, and 30 wt.%) and fungal species (*Ga* – *G. adspersum*; *Gs* – *G. sessile*). Data points outlined in black indicate composites that exhibited stress–strain curve shape II. Substrate abbreviations correspond to those listed in Table 1.

Slope 1 (A) represents the initial stiffness of the material and serves as an estimate of the compressive modulus. However, it should be noted that the actual elasticity of the samples was not determined. At a later stage of the compression test, Slope 2 (B in Fig. 3a) was measured. The compressive stiffness–density diagrams (Fig. 3b and 3c) reveal considerable variability in density and compressive stiffness across the different composite types, but consistent behavior among biological replicates. Composites bound with *G. sessile* appeared to have slightly higher densities than those with *G. adspersum*. This may be attributed to a thicker outer mycelium layer, which was particularly noticeable in composites produced with *G. adspersum*. In general, higher-density composites were stiffer, resulting in higher compressive moduli. Additionally, the grain spawn percentage appeared to have a slight influence on the modulus. Notably, composites made from pure birch bark exhibited the highest density and stiffness, likely due to the dense structure of birch bark, which contains numerous sclereids with thickened cell walls [9].

Interestingly, the differences between Slopes 1 and 2 are considerably greater in samples that exhibit high initial stiffness during the early stages of the compression test. This high stiffness is likely attributable to the compact arrangement of individual bark or wood particles, tightly held together by the ingrown mycelium. As the softer, weaker mycelium bond begins to yield or fracture, the particles start to shift relative to one another, resulting in a noticeable reduction in slope and overall stiffness. The compression test data allow for preliminary conclusions regarding the effectiveness of mycelium as a binder in bark composites. Large differences between Slopes 1 and 2 suggest that the mycelium is sufficiently robust to hold the bark particles in place, allowing the mechanical characteristics of the bark to influence the overall composite properties. In contrast, small differences between the two slopes indicate that the mycelium is less capable of forming strong connections between substrate particles. This behavior is particularly evident in composites made from Douglas fir bark and birch bark inoculated with *G. sessile* (data points outlined in black; Fig. 3b and 3c), and is also reflected in the characteristic shape of their stress–strain curves (curve type II; Fig. 3a). Despite the weak connection between substrate particles in these composites, the mycelium enabled the formation of stable bark–mycelium composites by developing a thick surface layer. The datapoints for ultimate compressive strengths (Fig. 4) support this hypothesis, since none of the composites was outstandingly weak. However, the ultimate compressive strength values were determined after the onset of permanent plastic deformation, most likely caused by exceeding the strength of the mycelium. Composites produced with *G. adspersum* developed a thicker surface mycelium layer but showed less internal colonization, resulting in lower density, greater flexibility, and lower compressive modulus values (Slope 1). In contrast, composites made with *G. sessile* exhibited more extensive internal mycelial growth and a thinner surface layer, which made them denser and stiffer. These two structural features—surface layer thickness and internal colonization (Fig. 5)—had a clear impact on the composites’ mechanical performance.

**Fig. 4:**
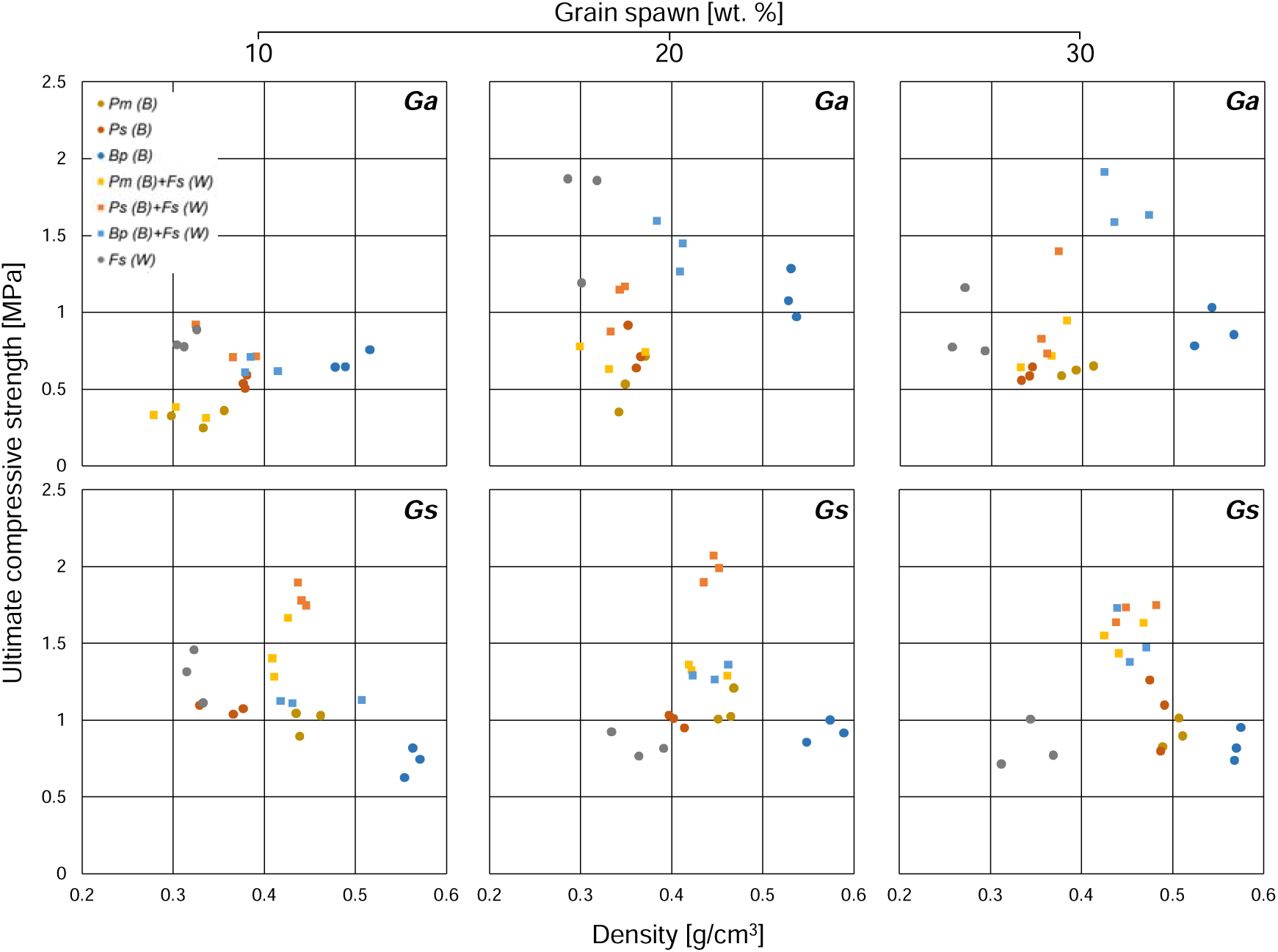
Ultimate compressive strength values derived from the region C illustrated in Figure 3a (maximum stress before failure) plotted against density. Data are presented separately for each grain spawn level (10, 20, and 30 wt.%) and fungal species (Ga – *G. adspersum*; Gs – *G. sessile*). Substrate abbreviations correspond to those listed in Table 1.

**Fig. 5:**
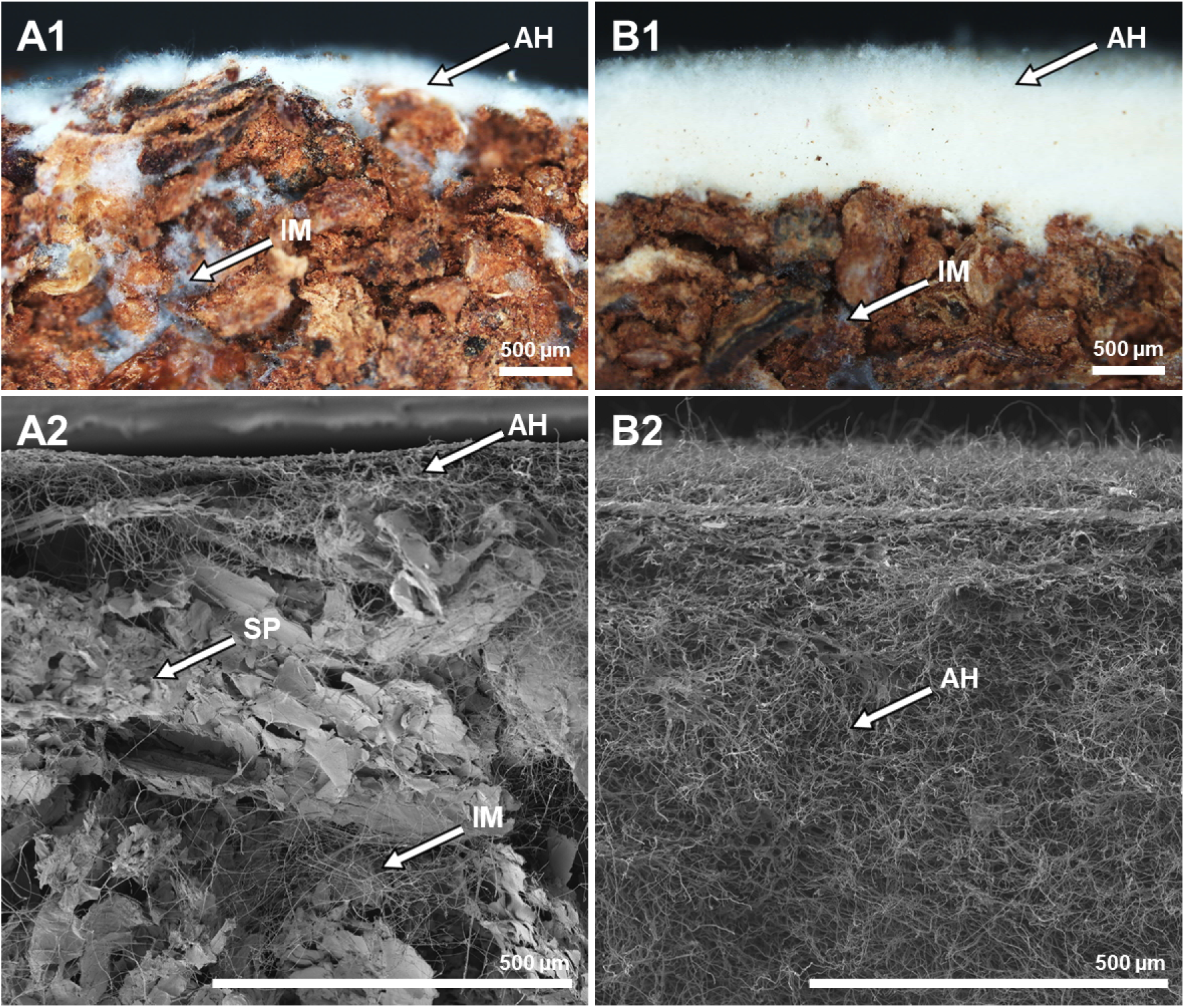
Cross-sectional images of mycelium-bound composites made of birch bark with 10 wt. % grain spawn produced for compression strength testing. Figures A1 and B1 show optical micrographs obtained using a binocular microscope, while A2 and B2 present corresponding environmental scanning electron microscopy (ESEM) images. (A1 and A2) *G. sessile* grown on birch bark (dense, stiff) forming a thinner but compact layer of aerial hyphae (AH) with more pronounced ingrown mycelium (IM) predominantly occupying interstitial spaces between substrate particles (SP). (B1 and B2) *G. adspersum* grown on birch bark (medium dense, flexible), showing a thicker, continuous layer of aerial hyphae (AH) with substantially less ingrown mycelium (IM). Scale bars: 500 µm.

The outer mycelium layer particularly strongly influenced the elastic response during compression. Composites with a pronounced outer layer behaved similarly to foam in the tests: they could withstand greater deformation before failure, and the mycelium surface eventually cracked under stress, as also described by [27]. This foam-like behavior was reflected in the Type II stress–strain curve, seen in composites made from Douglas fir bark (*Pm (B)*) and from a mixture of birch bark and beech wood (*Bp (B)* + *Fs (W)*), both inoculated with *G. sessile*.

### Water absorption potential

The water absorption potential of the produced composites was evaluated by immersing them in water for 72 hours. Four biological replicates were prepared for each substrate-fungus combination. Initially, the composites were removed from the water after 5 minutes, weighed, and re-immersed. While all composite variants exhibited a measurable increase in weight already within the first 5 minutes of immersion, the findings of this study showed that the overall water absorption potential is strongly dependent on the substrate-fungus combination (Fig. 6). The highest water absorption was observed for composites made from a mixed substrate of pine bark and beech wood with *G. adspersum* (224.7%) and in composites produced from pure bark with *G. sessile* (187.1%) after 72 hours. In general, composites produced with *G. adspersum* absorbed more water than those made with *G. sessile*. This difference is likely attributable to the thinner mycelium layer in *G. adspersum* composites, which increases open porosity and, consequently, water absorption (Fig. 7). In contrast, composites produced with *G. sessile* exhibited a thicker mycelium layer, reducing water absorption due to the hydrophobic properties of fungal proteins such as hydrophobins [37, 38]. In addition to the hydrophobic nature of mycelium, the properties of the lignocellulosic substrate also strongly influenced water absorption potential [39]. Composites produced with pure Douglas fir bark exhibited the lowest water absorption capacity for both fungal species. Conversely, composites produced with mixed substrates generally showed higher water absorption potential, likely due to the greater hydrophilicity of wood compared to bark.

**Fig. 6:**
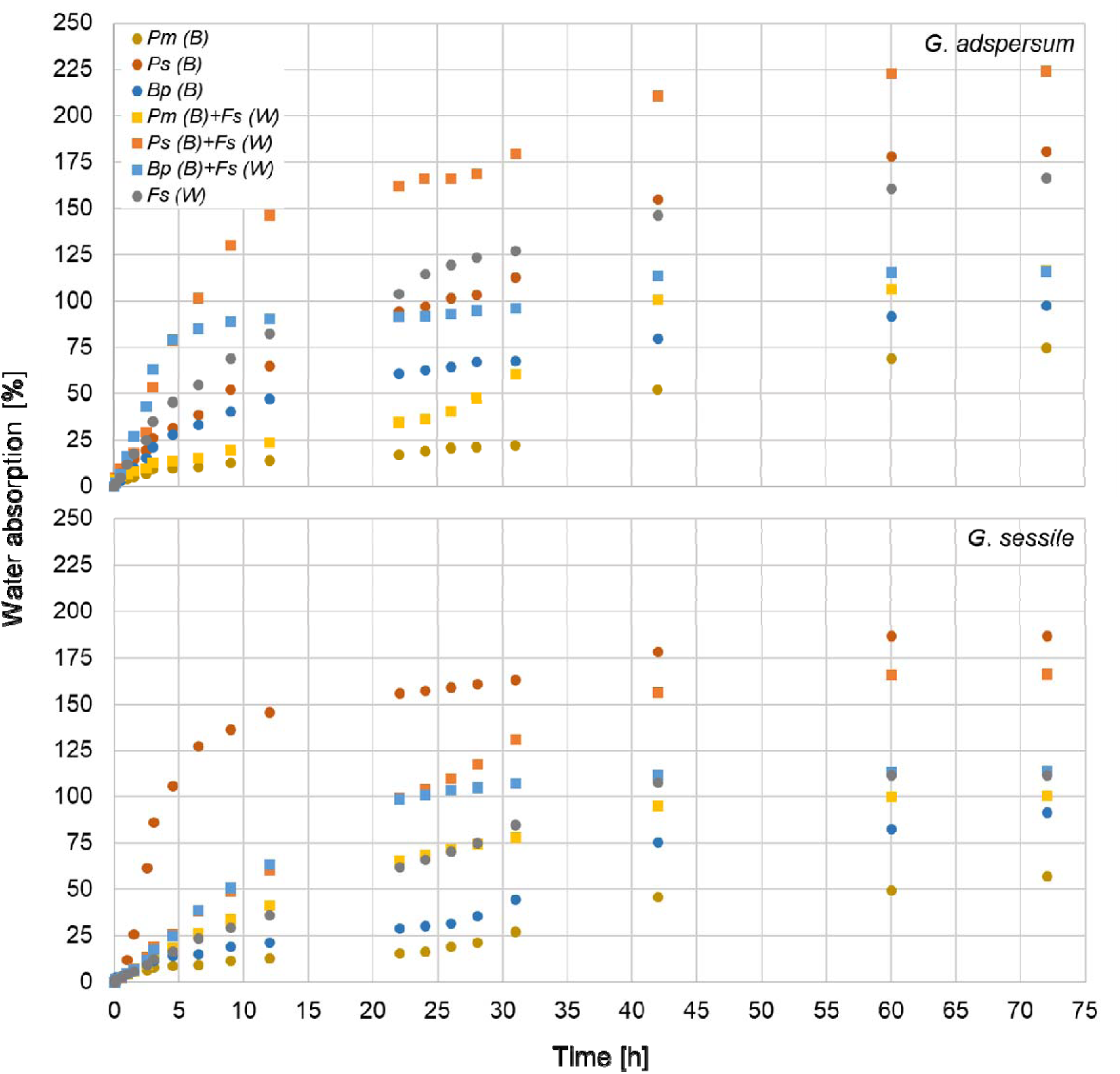
Water absorption (%) over time for all produced mycelium-bound composites. Each data point represents the mean value of 4 biological replicates (n = 4) produced only with 10% grain spawn.

**Fig. 7:**
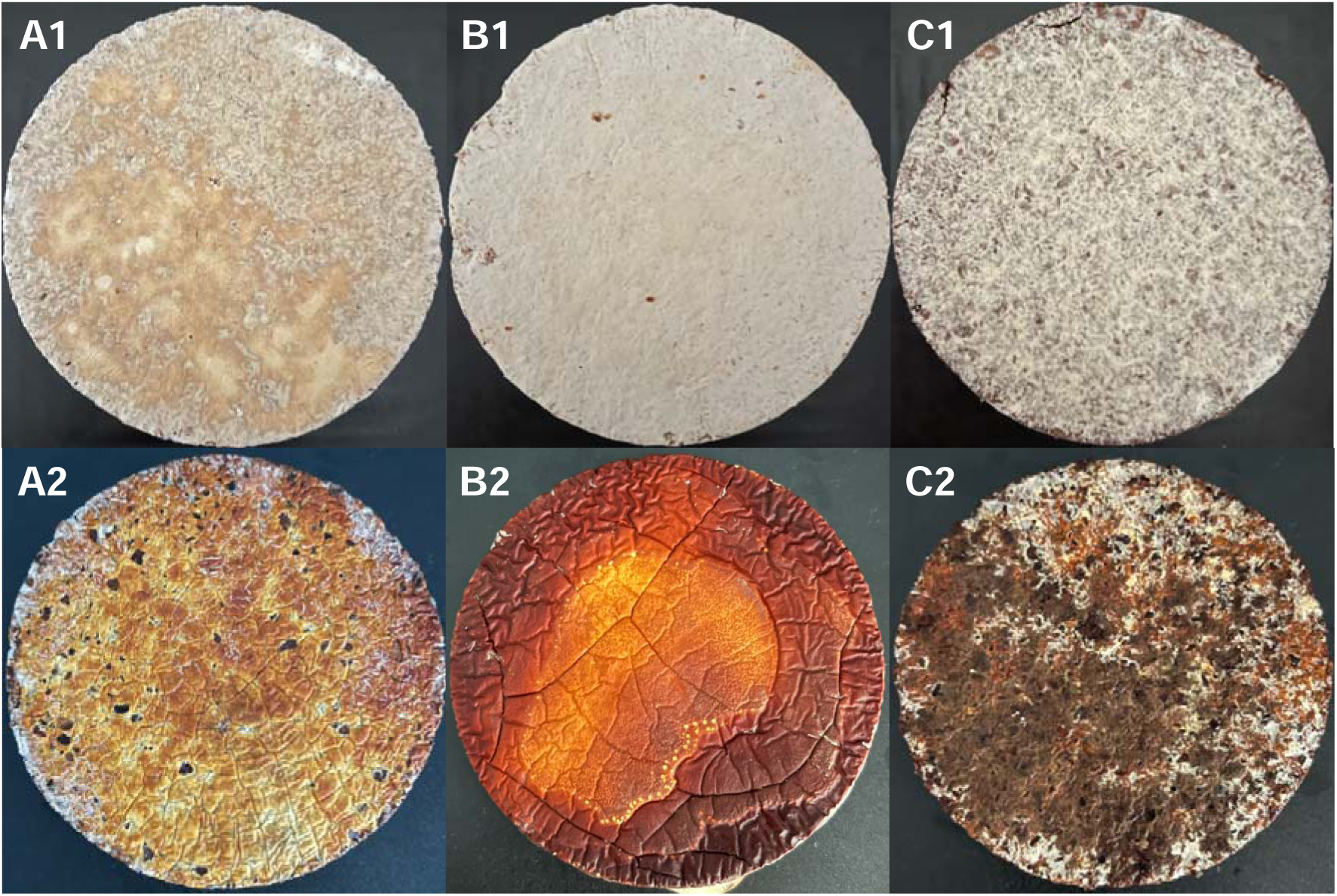
Images of the outer layer of mycelium-bound composites before (top row) and after 72 hours of water immersion (bottom row). (A) *G. adspersum* on pine bark with beech wood: thin mycelium layer with visible substrate particles and open spaces, allowing water infiltration. (B) *G. sessile* on Douglas fir bark with beech wood: very thick mycelium layer that restricts direct water entry, although the wood substrate remains hydrophilic. (C) *G. sessile* on pure Douglas fir bark: medium-thick mycelium layer with visible bark particles and a highly hydrophobic substrate.

### Morphological traits

Morphological analysis was conducted on all fungus–bark substrate combinations, as well as on beech wood inoculated with 10% grain spawn. Environmental scanning electron microscopy (ESEM) revealed consistent hyphal interactions with substrate particles across all samples. Observed colonization patterns included: (i) hyphae bridging voids between substrate particles to form an isotropic network [27] and (ii) hyphae overgrowing particle surfaces (Fig. 8). Although these morphological traits were common to all combinations, the extent of fungal colonization varied significantly with both fungal species and substrate type. Notably, bark-based substrates (I–III) exhibited fewer hyphae within void spaces compared to the beech wood substrate (IV), in which hyphae extensively colonized the interparticle regions. This variation in colonization density is likely to influence the microscale mechanical properties of the resulting mycelium-based composites. [2, 27].

**Fig. 8:**
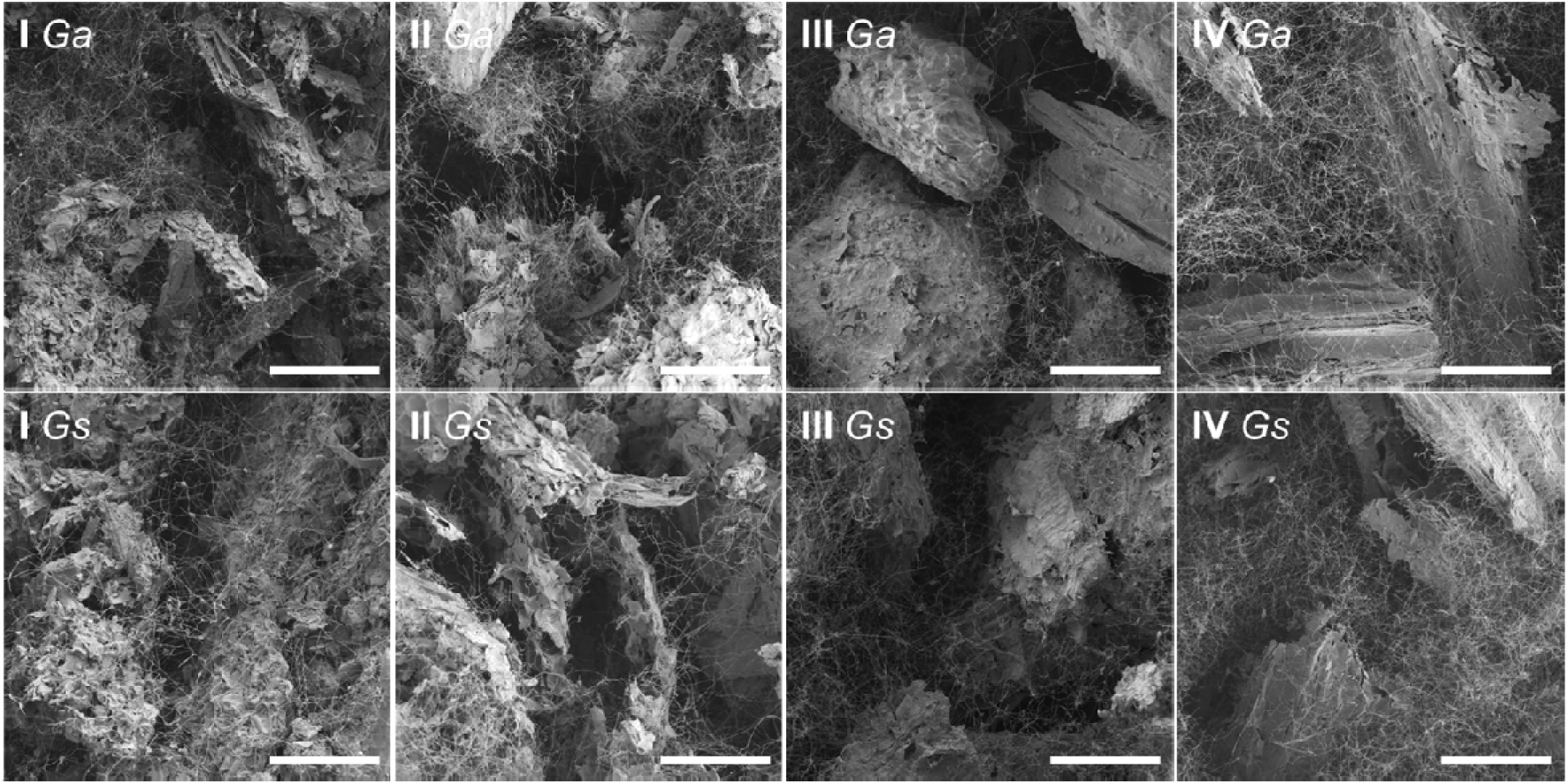
Environmental scanning electron microscopy (ESEM) images of *Ganoderma adspersum* (*Ga*) and *G. sessile* (*Gs*) colonizing substrates with following abbreviations: I – Douglas fir bark; II – Scots pine bark; III – European birch bark and IV – European beech wood. Scale bars: 150 µm.

### Conclusion

This study explored the feasibility of producing mycelium-bound composites using three types of bark (pure or mixed with beech wood) and two species of *Ganoderma*, under varying production conditions. A novel “growth-tube” method was developed to quantitatively evaluate fungal growth despite the high heterogeneity of bark and wood substrates. The results demonstrated that mycelium growth was primarily influenced by the type of substrate, with additional variation attributed to fungal species. Both fungi exhibited slower growth on bark substrates, likely due to their complex chemical composition and the presence of antifungal substances. Nevertheless, successful composite production was achieved across all fungus– substrate combinations, providing materials with diverse physical and mechanical properties. Composite density was influenced by the fungal species, substrate type, and the proportion of grain spawn (inoculum), ranging from 0.26–0.57 g/cm³ for *G. adspersum* and 0.31–0.59 g/cm³ for *G. sessile*. Composites made from mixed substrates generally exhibited lower densities, likely due to the lower density of wood particles and the development of a thicker mycelium layer, which increased the overall volume. A higher inoculum content was associated with increased composite density. Composites produced from pure bark substrates generally exhibited higher stiffness and brittle failure behavior, whereas those made from mixed substrates demonstrated greater flexibility, likely due to improved fungal colonization. In general, composites with *G. sessile* displayed higher strength and stiffness, making them suitable for structural applications. Conversely, *G. adspersum* composites showed enhanced flexibility, suggesting their potential for applications requiring impact absorption. Water absorption capacity was also strongly influenced by fungal colonization and substrate type. The highest absorption rates were observed in composites made from pine bark with beech wood and *G. adspersum* (224.7%), and from pure pine bark with *G. sessile* (187.1%). Composites produced with *G. adspersum* tended to absorb more water due to their thinner mycelium surface layer. In contrast, Douglas fir bark composites exhibited the lowest water absorption, likely due to the substrate’s hydrophobicity combined with pronounced mycelium layer.

Overall, the study confirms the potential of bark-based substrates for producing mycelium-bound composites using different fungal strains. It provides valuable insights into fungal growth behavior, composite morphology, and the resulting physical and mechanical performance, including water absorption characteristics. Future research should focus on enhancing fungal colonization through optimized fungus–substrate combinations, standardizing production processes, and improving composite properties for specific applications as sustainable alternatives to conventional materials.

## Notes

### Competing Interest Statement

The authors have declared no competing interest.

### Funding

- Federal Ministry of Education and Research: 033L303B

